# A Novel Method for Normalizing Data from DNA-Encoded Library Selections

**DOI:** 10.64898/2026.01.20.700605

**Authors:** Zsofia Lengyel-Zhand, Zhaowei Jiang, Justin I. Montgomery, Hongyao Zhu, Keith Riccardi, Richard Corpina, Woodrow Burchett, Mario Abdelmessih, Robert Stanton, Timothy K. Craig, Timothy L. Foley

**Author notes:** These authors contributed equally (Zsofia Lengyel-Zhand, Zhaowei Jiang).

## Abstract

DNA-encoded library screening represents a significant advancement in the field of drug discovery. Its ability to rapidly and cost-effectively identify potential drug candidates from large compound libraries has the potential to revolutionize the way new medicines are discovered and developed. While the strategies for DEL screening and data analysis have improved over the years, data normalization remains an open challenge. Existing normalization methods can yield poor correlation for compounds with high read count, and they do not account for inherent sources of noise. To overcome these drawbacks, we have developed a robust normalization technique using an antibody fragment and a DNA-conjugated peptide as an internal control. This innovative approach allows for normalization between samples of different conditions and accounts for technical challenges that occur during screening.

**Table of Contents Graphic:** 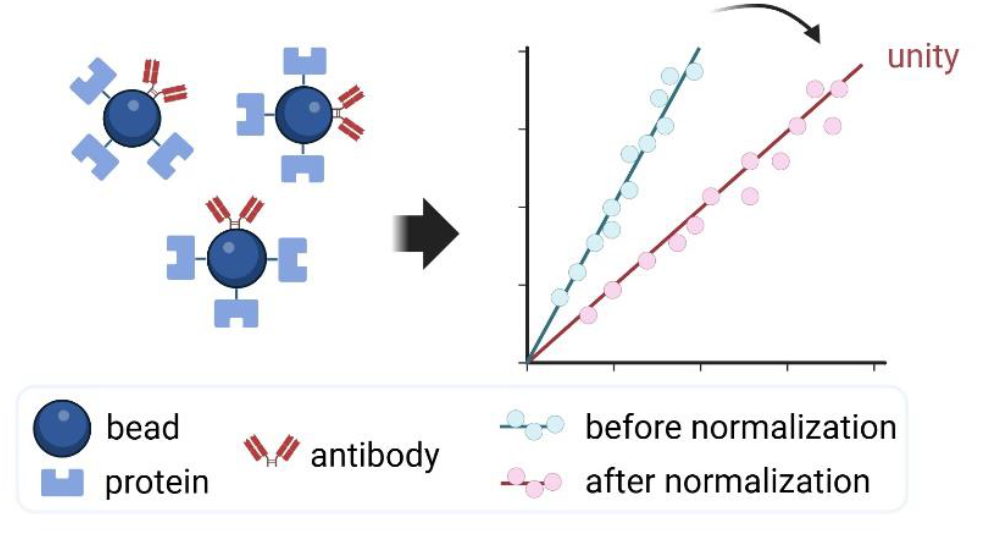

**Synopsis:** Normalization of DNA-encoded library selection data reduces bias and noise, enabling accurate identification of true binders and reliable enrichment analysis.

## 1. Introduction

DNA-encoded library (DEL) screening has emerged as a transformative technology in early-stage drug discovery, enabling rapid identification of small molecule binders to biological targets.^1-3^ It allows for screening of enormous compound libraries, where each compound is covalently tagged with a unique DNA sequence that functions as a molecular barcode. The screening process typically involves multiple rounds of affinity selections, where DEL compounds are incubated with a target protein immobilized on a surface, followed by PCR amplification and next-generation sequencing of the enriched barcoded DNA.

The impact of DEL screening on drug discovery has been profound.^3-7^ It has enabled the exploration of vast chemical spaces, allowing for the discovery of novel compounds with therapeutic potential.^8-12^ It is often useful to compare binding of specific chemical matter under different screening conditions, for example target vs no target control, with or without known ligands, with mutated targets, with selectivity targets and so forth, and these comparisons can offer insight into the molecular mechanisms through which these ligands operate. Additionally, DEL screening can lower drug discovery costs by enabling the testing of billions of compounds in a single experiment, reducing the resources needed compared to traditional high-throughput screening, while expanding the diversity of chemical matter accessible to researchers.^6,13^ Although the methods for DEL compound synthesis and screening have undergone significant advancements over time,^14-19^ data analysis and reliable hit identification can still be challenging. Variability in sequencing depth, compound representation, and experimental conditions can obscure true binders and complicate downstream analysis. Directly comparing the sequencing read counts from next generation sequencing of the DEL barcodes does not provide a good approach for comparing data between conditions, and some form of normalization of read counts is required to allow quantitative assessment. To date, there is no consensus on how to normalize read counts across samples within an experiment and there are only a few published methods that deal with noisy data and normalization techniques. Kuai et al. proposed a normalization workflow based on a published protocol, that compares the observed read count of each library member to an expected value, which is calculated from the ratio of the total diversity of the library and the total number of reads.^20^ This approach assumes that library members are evenly distributed prior to selection, even though the actual distribution of members in any DEL library is unknown. Conversely, the method developed by the Schreiber lab accounts for the low abundance of compounds that were determined experimentally.^21^ Their model calculates a “normalized fold change” score by employing Poisson distribution based on the count of observed sequencing reads.

Building on these foundational methods, recent advancements in artificial intelligence (AI) and machine learning (ML) have introduced powerful tools for denoising, hit ranking and normalizing DEL selection outputs.^22-26^ A notable advancement is the sparse learning-based method “deldenoiser”, which models DEL data to recover meaningful hits in the presence of noise and synthetic artifacts.^23^ As DEL screens often suffer from confounding variables such as bead binder and protein immobilization artifacts, deldenoiser uses patterns common across cycles and compounds to suppress these confounding signals. This method significantly improves recovery of true binders and better reflects the underlying structure-activity relationships compared to naïve count-based ranking. Moreover, McCloskey et. al applied ML to DEL data to improve hit finding by filtering out noise and non-specific binders.^22^ Their model predicted a potent, novel compound beyond traditional enrichment analysis, identifying a hit structurally distinct from both the DEL library and known ligands. Recently, Iqbal et al.^26^, conducted a comparative study evaluating different combinations for DEL screening and ML models for hit discovery. Using three DELs and multiple ML methods, their key finding indicated that although model accuracy varied, 10% of predicted binders and 94% of non-binders were confirmed experimentally. This study emphasized the importance of training data diversity and ML model generalizability rather than relying solely on prediction accuracy.

While offering good noise handling and scalability, ML methods require careful model design, reliable training data, and validation to avoid false positives or biased outputs. Additionally, machine learning has not proven effective for normalizing DEL data across replicates or screening conditions, which remains a critical challenge in DEL analysis. A successful normalization method across replicates should return the same enrichment value for a given DEL compound. As a result, the read count plot has symmetrical patterns and enrichments scatter around a unity line (**Figure 1**.). The Pfizer DEL group had previously investigated data noise and uncertainty in DEL selections by analyzing replicate datasets.^27^ They introduced a method using logarithmic transformation of read counts and normalization across replicates to estimate experimental noise. Their results showed that noise levels correlate strongly with sequencing depth and compound abundance. Notably, they demonstrated that removing compounds with low read counts (1-5 reads) significantly reduces noise and improves hit reliability – boosting confirmation rates by more than 100-fold for higher-count subjects. Although this method operates independently of selection conditions and offers a broad framework for enhancing DEL data quality, it does not effectively support normalization across different samples.

**Figure 1.**
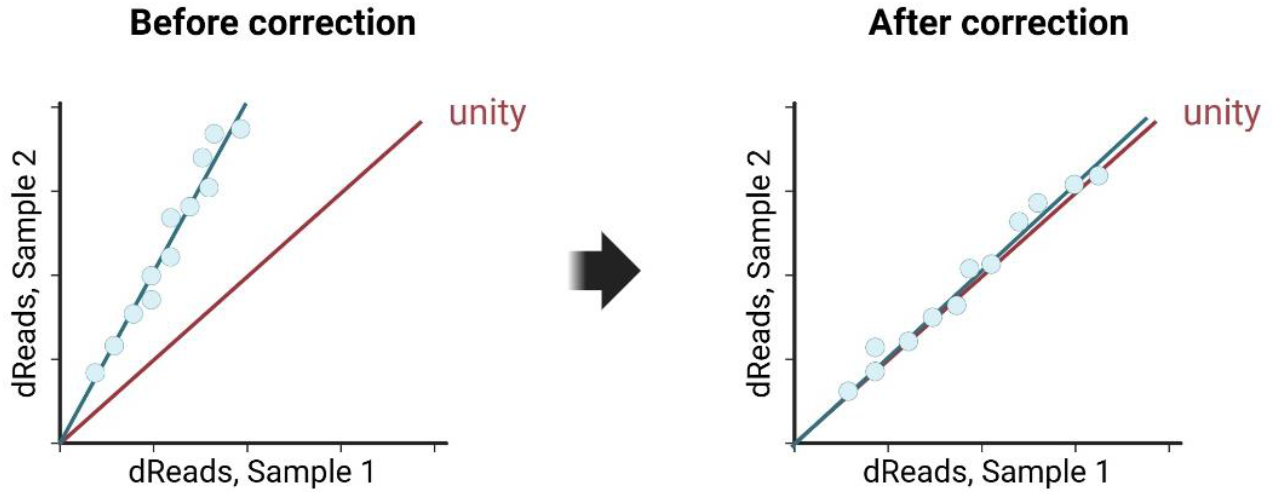
Graphic illustration demonstrates counts between replicate samples before (left) and after (right) normalization. Prior to correction, systematic bias causes deviations from the expected unity line.

Here, we introduce a new normalization method that utilizes an anti-hemagglutinin (anti-HA) antibody and an on-DNA HA-peptide pair. By adding anti-HA antibody to all DEL samples and spiking DNA-conjugated HA-peptide into the DEL pool, we can assess peptide enrichment across different samples and use this data to adjust the dRead values of the DEL compounds (**Figure 2**.). We show that this approach enables effective normalization not only between biological replicates but also among samples subjected to varying screening conditions, thereby improving the reliability and comparability of the results.

**Figure 2.**
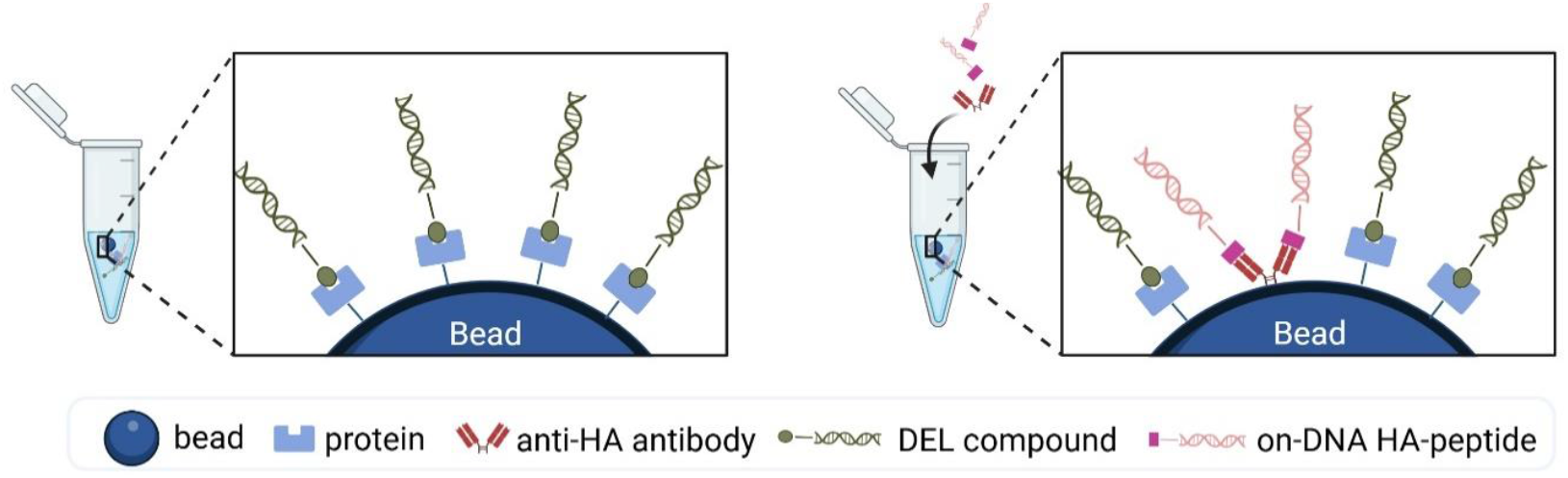
Schematic illustration of spike-in control strategy of DEL screening. In a typical DEL screening workflow, target proteins are incubated with the DEL pool and subsequently immobilized on beads (left panel). To facilitate cross-sample normalization and minimize assay variability, a spike-in control is introduced during the binding assay (right panel). This system includes an anti-HA antibody and an HA-peptide conjugated to DNA, serving as internal controls to ensure consistent performance and reliable data interpretation.

## 2. Results and Discussion

Accurate interpretation of DEL screening data across various samples remains a key challenge due to inherent noise and variability introduced during selection, amplification, and sequencing. To address this problem, we chose to include an anti-HA antibody/HA-peptide pair in DEL selections as an internal control. This allows us to achieve normalization among samples subjected to various screening conditions, ensuring more reliable and comparable results.

### 2.1 Validation of antibody for DEL screening

To determine whether an antibody could serve as an internal control for DEL normalization, we examined its performance across two key criteria: binding specificity and enrichment capacity. We assessed its binding function after immobilization using the bead-assisted ligand isolation fluorescence assay (BALI-FL), which was originally developed by Foley et al.^29^ This assay can provide a quantitative measure of the active site content of the reagent after immobilization. In the experiment, the antibody was incubated with increasing concentrations of fluorescently labeled HA-peptide, then captured, along with the bound ligands, using magnetic affinity beads to enable separation from the unbound fraction. The bound ligand was eluted, and both bound and unbound fractions were analyzed with a fluorescence plate reader. As illustrated in **Figure 3a**, there was a strong accumulation of tracer in the bound fraction up to ∼1000 nM peptide, indicating effective and saturable binding. At the same time, only minimal tracer was found in the supernatant, which supports the presence of high-affinity interaction between the antibody and the peptide. Taken together, these findings confirm that the immobilized anti-HA antibody maintains its binding capacity after immobilization and is appropriate for use in subsequent DEL selections.

**Figure 3.**
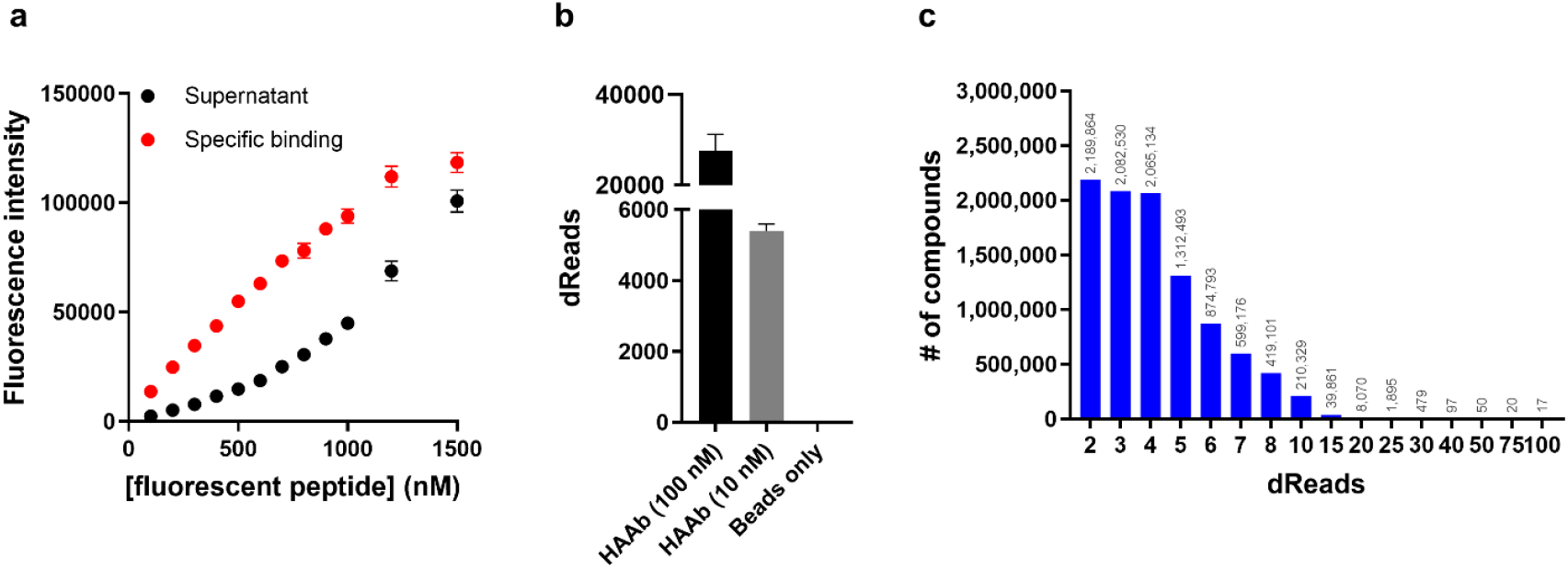
Anti-HA antibody shows high enrichment of HA-peptide. (a) antibody retains its binding capacity after immobilization as shown by BALI-FL. Increasing concentration of fluorescent tracer was titrated with a constant input antibody concentration of 500 nM. Clear inflection points were observed in the supernatant and eluate (specific binding) fraction that indicates the stoichiometric titration of binding sites within the sample. (b) anti-HA antibody showed enrichment of the on-DNA peptide after 2 rounds of DEL selection. (c) number of compounds vs dRead values after DEL screening. The antibody did not enrich DEL compounds as fewer than 500 compounds showed more than 30 dReads and those had been tagged as frequent hitters.

Next, we evaluated the antibody’s ability to enrich DNA-conjugated HA-peptide in a DEL selection. The anti-HA antibody was subjected to 2 rounds of DEL selection, using Pfizer’s proprietary DNA-encoded library pool spiked with the on-DNA HA-peptide. Parallel screening conditions were pursued, including various target concentrations and a beads only control. The DEL screening showed that the anti-HA antibody effectively enriched the HA-peptide even at low target concentrations (10 nM), while the beads only control showed negligible signals (**Figure 3b**). This demonstrates the antibody’s high specificity and sensitivity. Notably, the antibody did not enrich other DEL compounds, as fewer than 500 compounds accumulated more than 30 dReads (**Figure 3c**), and those compounds were already identified as bead binders or frequent hitters in previous DEL selections. Overall, these results indicate that the antibody performs reliably in DEL selections, effectively enriching its peptide ligand without causing any confounding DEL enrichment artifacts.

### 2.2 Validation of antibody/peptide pair as internal controls in DEL selections

Our normalization strategy was tested through several DEL selection experiments, involving protein targets from multiple target classes. In each DEL screen, 2.5 μM of protein and 100 nM anti-HA antibody were incubated with Pfizer’s DEL pool consisting of 35 libraries and 1,000,000 copies of the DNA-tagged HA-peptide, the same number of copies as each individual library member. In each DEL selection, target samples had a biological duplicate (Samples 1 vs 2 as shown in **Figure 4**). Following DEL selection and next-generation sequencing, the dRead count for the HA-peptide were used as a normalization reference to enable comparison across independent samples. For each sample, a normalization factor (**Equation 1**) was determined by dividing the HA-peptide read count of the specific sample by the highest HA-peptide read count observed in that screening set. The dRead values for the DNA tagged HA-peptide and the calculated normalization factors for all the samples in the different DEL screenings are summarized in **Table S1**. in the Supplementary Information. These normalization factors were then applied to adjust the DEL dRead values, thereby compensating for technical variability between samples (**Equation 2**).

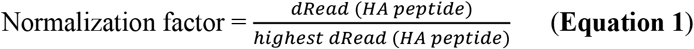

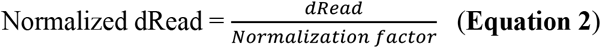

**Figure 4.**
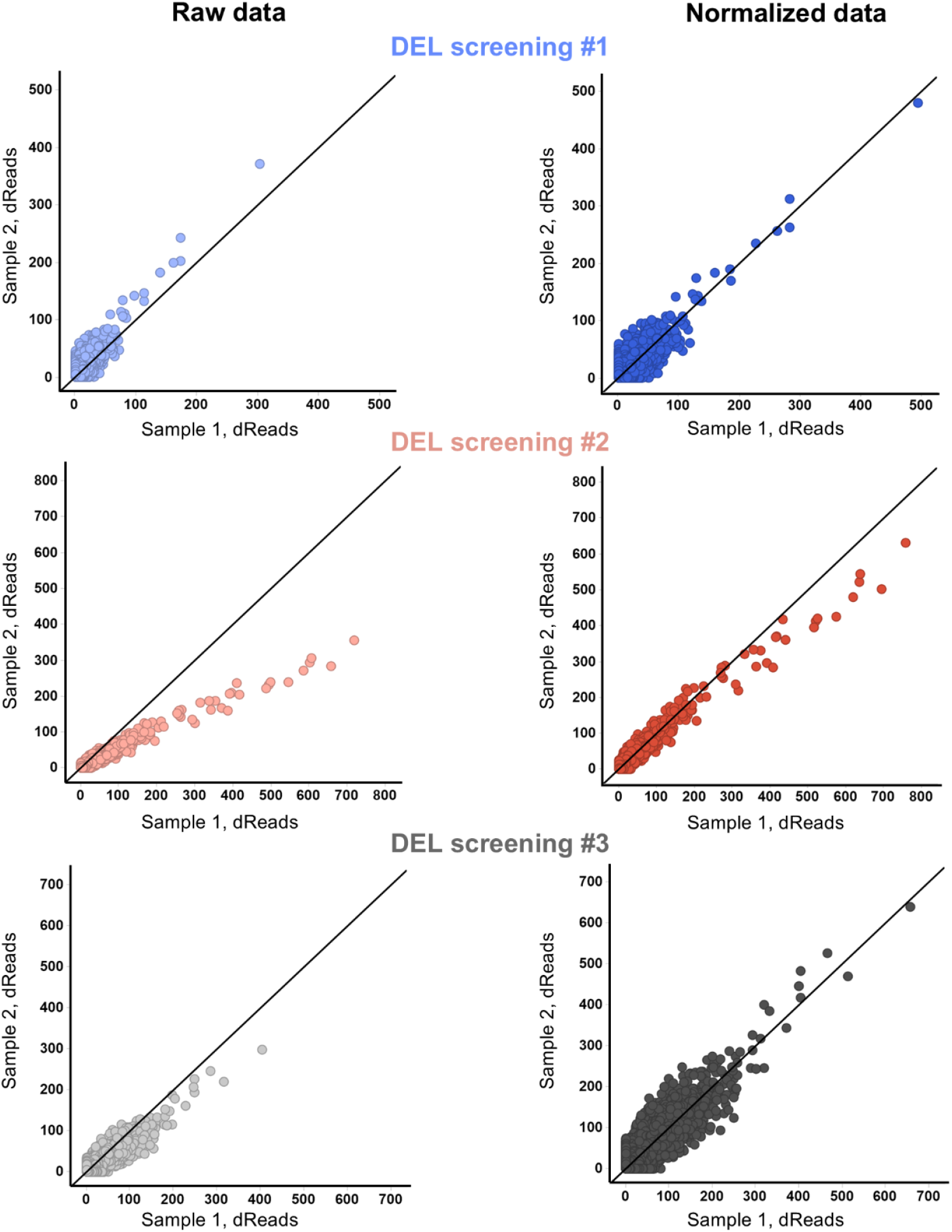
Normalization successfully returns read counts to unity in all three DEL screenings. The read count scatter plots compare raw data with normalized data for 3 different DEL campaigns (3 different target proteins). For all campaigns, sample 1 vs 2 are biological replicates and their enrichment should scatter around the unity line, however due to systemic errors, raw read counts shift away from the diagonal line, favoring either sample 1 or 2. After applying our normalization method, read counts are returned to unity, indicating successful normalization.

A reliable normalization approach for biological replicates should produce consistent enrichment values for each DEL compound, leading to symmetrical patterns in the read count plot and clustering of enrichment values near the unity line. As shown in **Figure 4**, the raw data for Samples 1 vs 2 (biological duplicates in each DEL selection) initially deviates from the unity line in all 3 DEL selection campaigns. However, after applying our normalization technique, the dRead values align closely with the unity line, indicating improved consistency across replicates.

To assess how data normalization affects the interpretation of target selectivity, we analyzed the enrichment of compound 1 in a DEL selection targeting kinase proteins. In biochemical assays, compound 1 previously demonstrated an IC_50_ of 135 nM for Kinase 1 (K1) and 873 nM for Kinase 2 (K2), indicating a stronger affinity for K1. However, the raw DEL sequencing data indicated similar enrichment of the compound for K2 (705 dReads) and K1 (471 dReads), which did not accurately reflect the compound’s selectivity. After applying our internal antibody control to normalize the data across samples, the revised read counts showed a notable shift, with K1 displaying significantly higher enrichment (2944 reads) compared to K2 (892 reads) (**Figure 5**). This adjustment aligned the DEL results with those from the biochemical assays, highlighting the critical role of normalization in reliably identifying selective binders during multi-target screening efforts.

**Figure 5.**
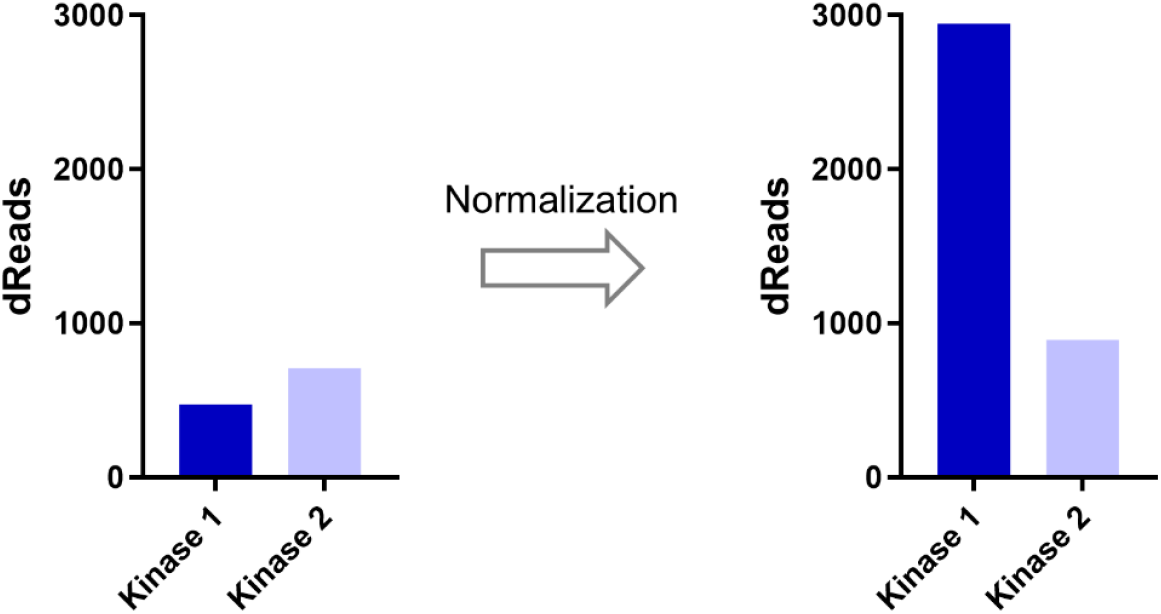
Normalization improves comparability of DEL enrichments. Raw read counts for Kinase 1 and Kinase 2 initially suggest comparable enrichment levels, implying similar affinity for the compound of interest. However, following normalization, the dRead for Kinase 1 show a marked increase, suggesting a higher affinity for the compound. This demonstrates how normalization can uncover meaningful differences that are obscured in raw data, thereby improving the interpretability and reliability of DEL screening results.

### 2.3 Engineering Antibody Fragments for normalizing DEL selections

Although the full-length anti-HA antibody is highly effective as an internal control, it has some limitations. The antibody is expensive, producing it in-house would be challenging, and its biotin label makes it incompatible with DEL selections involving target proteins with alternate affinity tagging systems. To overcome these challenges, we designed a single-chain variable fragment (scFvs) of the anti-HA antibody (**Figure S1**). Antibody fragments provide several benefits, such as a smaller size, the capability for internal expression and purification, and the option to add different affinity tags to either the N- or C-terminal of the protein.

For DEL applications, reagents must remain stable under diverse conditions, including the presence of reducing agents (e.g., DTT, TCEP) that can compromise conventional antibodies by disrupting disulfide bonds. To address these concerns, we identified an intrabody from the literature that recognizes the HA epitope inside living cells^30^, where the reducing environment normally renders most nanobodies non-functional. We cloned this nanobody with C-terminal tags to pET28a and co-expressed the construct in *E. coli* with an additional expression plasmid for *E. coli* BirA. The protein was purified via Ni-NTA affinity chromatography followed by size exclusion chromatography. SDS-PAGE analysis (**Figure S2a**) and mass spectrometry spectra (**Figure S2b**) verified the successful expression of the biotinylated scFv.

The purified anti-HA scFv was validated in a similar way as the antibody. The available binding sites of the immobilized scFv were assessed using BALI-FL.^29^ At a constant scFv concentration (500 nM), peptide titration showed clear dose-dependent binding, with significantly higher fluorescence in the bound fraction compared to the supernatant (unbound), confirming that the active binding site remained accessible after immobilization (**Figure S3**).

We also evaluated the scFv in DEL selection to confirm its ability to specifically enrich the DNA-conjugated HA-peptide. The scFv was immobilized on neutravidin magnetic beads and following two rounds of DEL selection and sample sequencing, we detected high enrichment of the peptide at different scFv target concentrations (**Figure S4**). This result confirms that the designed scFv is able to enrich the DNA-conjugated HA-peptide and is appropriate for use as an internal control in DEL selections. Future studies will aim to assess its application in DEL selections with target proteins that have tags other than biotin.

## 3. Conclusion

DEL screening has revolutionized drug discovery by allowing rapid and cost-effective identification of drug candidates from large compound collections.^6^ While there have been improvements in DEL synthesis, screening, and data analysis, normalizing sequencing data remains a challenge, especially when comparing samples under different conditions. Current normalization methods often do not adequately address technical noise and variability, particularly for high read count compounds. This study presents a new normalization strategy that uses an anti-HA antibody and a DNA-linked HA-peptide pair as internal controls. This method enables normalization across biological replicates and different screening conditions, increasing the reliability and clarity of DEL data. Validation experiments showed the antibody’s specific binding and enrichment ability, and its effectiveness in resolving inconsistencies between raw DEL data and biochemical assays. Additionally, engineered scFv fragments of the antibody provide a cost-effective and flexible alternative for normalization, offering low background and high specificity in DEL selections. By filling a key gap in DEL data analysis, this work introduces a valuable tool to enhance consistency and confidence in drug discovery using DEL technology.

## Acknowledgments

The authors wish to acknowledge Sylvie K Sakata, Anthony R Harris, Shi Chen and all members of the Pfizer DEL team and HitGen team for their support.

