## Supporting figure S1 for "A Novel Method for Normalizing Data from DNA-Encoded Library Selections"

**Materials**

DELs were prepared using a split-and-pool methodology at HitGen Ltd. (Chengdu, China) following a general strategy similar to that described in Kung et al.[1] Neutravidin SpeedBeads (GE Healthcare, Piscataway, NJ) were used in this study. Mouse Biotin anti-HA.11 tag, 16B12 was purchased from Biolegend (cat#: 901505, San Diego, CA). 4-(2-hydroxyethyl) piperazine-1-ethanesulfonic acid sodium salt (HEPESNa) and 5M sodium chloride (NaCl) were purchased from Thermo Fisher Scientific (Waltham, MA).

**Buffers**

| **Buffer Name** | **Abbreviation** | **Recipe** |
|---|---|---|
| Bead-assisted ligand isolation buffer | BALI buffer | 50mM HEPESNa<br>150mM NaCl<br>0.01% (v/v) Tween 20<br>pH 7.4 |
| Antibody fragment elution buffer | Buffer AE | 20 mM Tris pH 8.0<br>0.2 M NaCl<br>10% (v/v) Glycerol |
| DEL selection buffer | Buffer DS | 50 mM HEPESNa<br>150 mM NaCl<br>0.3 mg/mL ssDNA<br>0.01% (v/v) Tween 20<br>pH 7.4 |
| DEL wash buffer | Buffer DW | 50 mM HEPESNa<br>150 mM NaCl<br>0.3 mg/ml ssDNA<br>0.01% (v/v) Tween 20<br>pH 7.4 |
| DEL elution buffer | Buffer DE | 50 mM HEPESNa<br>150 mM NaCl<br>0.01% (v/v) Tween 20<br>pH 7.4 |

### Methods

#### Bead-Assisted Ligand Isolation

Bead-assisted ligand isolation fluorescence assay (BALI-FL) was performed via 500 nM antibody or antibody fragment incubation with an increasing concentration Cy5-conjugated HA-peptide for 1-hour at room temperature (RT) in a total volume of 40 µL BALI buffer. Following this equilibration step, the samples were transferred to wells of a PCR plate that contained prewashed Neutravidin Speedbeads. A "peptide only" sample was carried through in parallel for use as control to assess for bead binding of the peptide ligand. The beads were resuspended and allowed to be incubated at RT for 30 min. Magnetic separation was performed with a ring magnet. The supernatant was aspirated to a receiver plate, and the peptide bound to the antibody was eluted by heating the sample at 95°C for 20 min on a thermocycler. Following transfer of the eluate to the receiver plate, the fluorescence of fractions was detected on a PHERAstar plate reader (BMG labtech, Cary, NC).

#### Antibody fragment design and purification

The anti-HA scFv franken body XHA15F11 sequence was adapted from Zhao et. al.,[2] and a linker, Avi-tag, structured linker, was appended to the N- or C-terminus. Sequences were codon optimized for *Escherichia coli* expression and cloned into pET28a at Genscript (Piscataway, NJ). Sequence of the protein is provided as a genbank file. Plasmid was co-transformed into *E. coli* BL21DE3 produced in-house with an additional expression plasmid for *E. coli* BirA, and protein expression was performed in Magic Media at 20°C for 48 hours. Proteins were purified by Ni-NTA affinity chromatography and size exclusion chromatography into Buffer AE.

### DEL selection and data analysis

Pfizer DELs were used in the selection and automated selection was carried out using a KingFisher Duo Prime Purification System (ThermoFisher, Waltham, MA) in 96-deep well plate. 2.5 µM of protein of interest and 100 nM anti-HA antibody were incubated with Pfizer DEL pool, containing the DNA-conjugated HA-peptide, in Buffer DS at RT for 1 h, followed by 30 min incubation with Streptavidin Magnetic Beads. Immobilized protein was washed for 1 min at RT in 200 µl Buffer DW 4 times. Retained DEL members recovered by heat elution in Buffer DE at 95 °C for 20 min. After the first round of selection, second round was repeated with the eluted portion of previous round used as the input to the successive round with protein/antibody. After each round, the output was quantified by qPCR. After two rounds, the selection was done and the output was amplified by PCR, then sequencing was performed for PCR amplified samples on an Illumina NextSeq2000 sequencer (Illumina Inc., San Diego, CA). All sequences were decoded by mapping a specific region of the observed DNA sequence back to the library ID and corresponding building blocks. The number of times each unique DNA barcode is seen is referred to as "count". After decoding, the reads which were duplicates due to PCR amplification were excluded through a pivot on a degenerate code which was randomly generated as part of the tag on each compound. In this study, pivoted sequence counts are referred to dReads, which represent molecule counts of individual chemical structures of DELs identified by their corresponding DNA tags.[3]

**Supporting Table**

Supporting Table S1

| DEL screening #1 | | |
|---|---|---|
| **Sample ID** | **On-DNA HA-peptide dRead** | **Normalization Factor** |
| **1_1** | 1505 | 0.613 |
| **1_2** | 1900 | 0.774 |
| **1_3** | 2007 | 0.818 |
| **1_4** | 2455 | 1.000 |
| **1_5** | 2111 | 0.860 |
| **1_6** | 2301 | 0.937 |
| **1_7** | 2361 | 0.962 |
| **1_8** | 1655 | 0.674 |
| **DEL screening #2** | | |
| **Sample ID** | **On-DNA HA-peptide dRead** | **Normalization Factor** |
| **2_1** | 651 | 0.948 |
| **2_2** | 388 | 0.565 |
| **2_3** | 414 | 0.603 |
| **2_4** | 442 | 0.643 |
| **2_5** | 552 | 0.803 |
| **2_6** | 687 | 1 |
| **2_7** | 538 | 0.783 |
| **2_8** | 474 | 0.690 |
| **DEL screening #3** | | |
| **Sample ID** | **On-DNA HA-peptide dRead** | **Normalization Factor** |
| **3_1** | 11482 | 0.615 |
| **3_2** | 8728 | 0.467 |
| **3_3** | 6558 | 0.351 |
| **3_4** | 9655 | 0.517 |
| **3_5** | 17153 | 0.918 |
| **3_6** | 18679 | 1 |
| **3_7** | 14103 | 0.755 |
| **3_8** | 10683 | 0.572 |

**Supporting figures**

Supporting figure S1

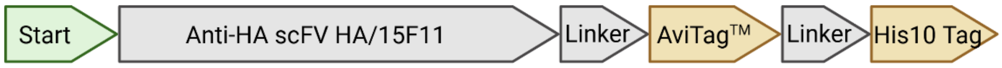

**Figure S1.** Schematic representation of the Anti-HA scFv HA/15F11 construct.

Supporting figure S2

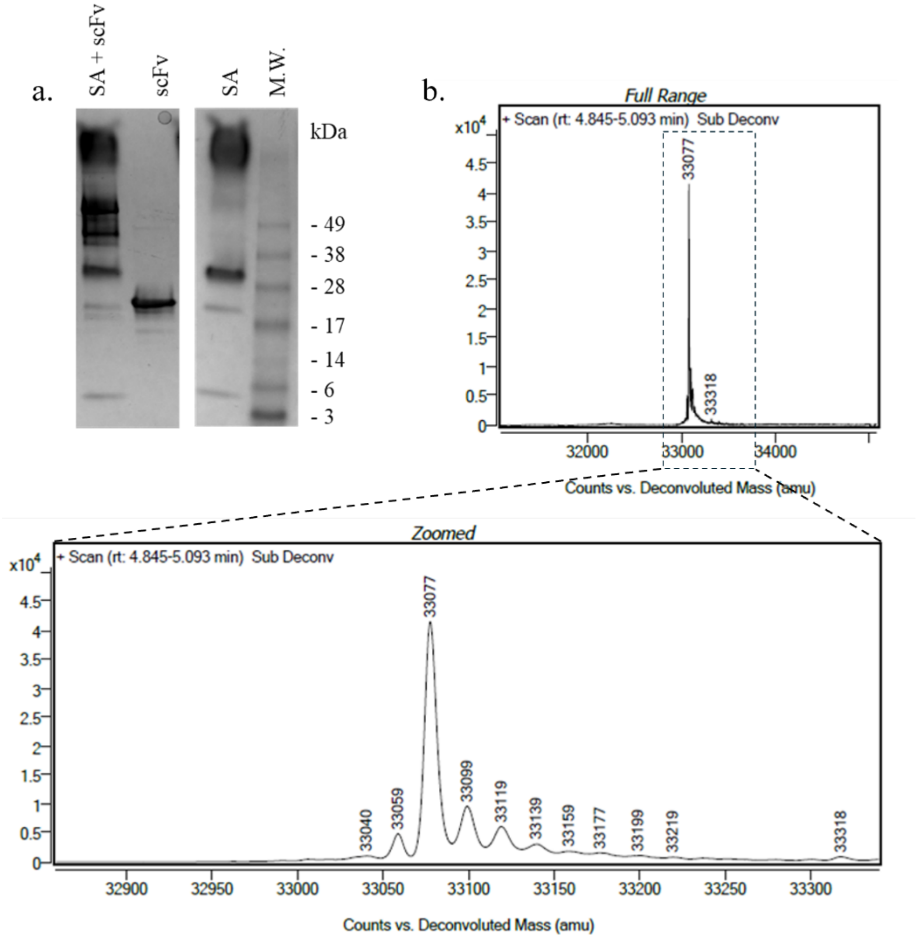

**Figure S2.** Characterization of purified scFv. (a) SDS-PAGE analysis confirms complex formation between scFv and streptavidin, confirming that the purified protein is biotinylated. (b) deconvoluted mass spectrometry spectra showed the expected biotinylated scFv mass, with zoomed-in view resolving the scFv peaks.

Supporting figure S3

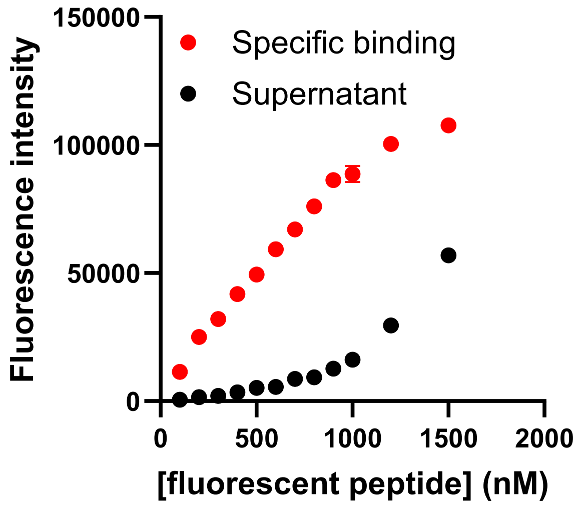

**Figure S3**. BALI-FL validation with fluorescent peptide titrated with a constant input scFV concentration of 500 nM. scFv were immobilized with Neutravidin beads.

Supporting figure S4

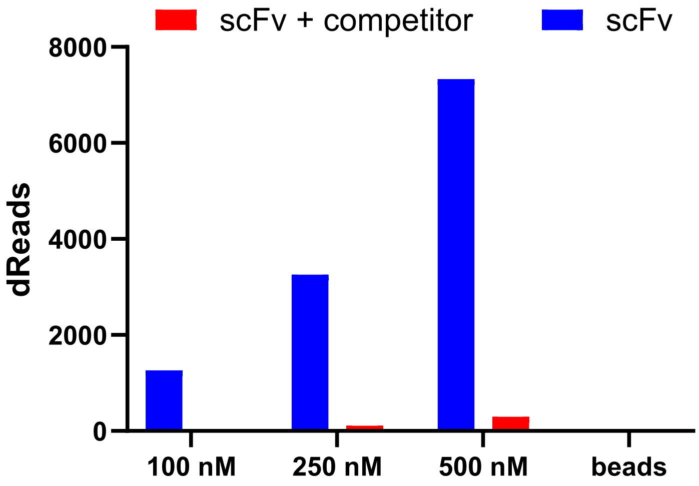

**Figure S4.** DEL screening results for scFv and its corresponding controls. After 2 rounds of DEL selection, scFv shows clear peptide enrichment even at the lowest (100 nM) concentration. Binding of the peptide was effectively blocked by a competitor (high concentration of HA peptide without DNA tag), suggesting specific binding of the DNA-conjugated peptide to the scFv.
